## Supplemental data for "Comparative therapeutic strategies for preventing aortic rupture in a mouse model of vascular Ehlers Danlos syndrome"

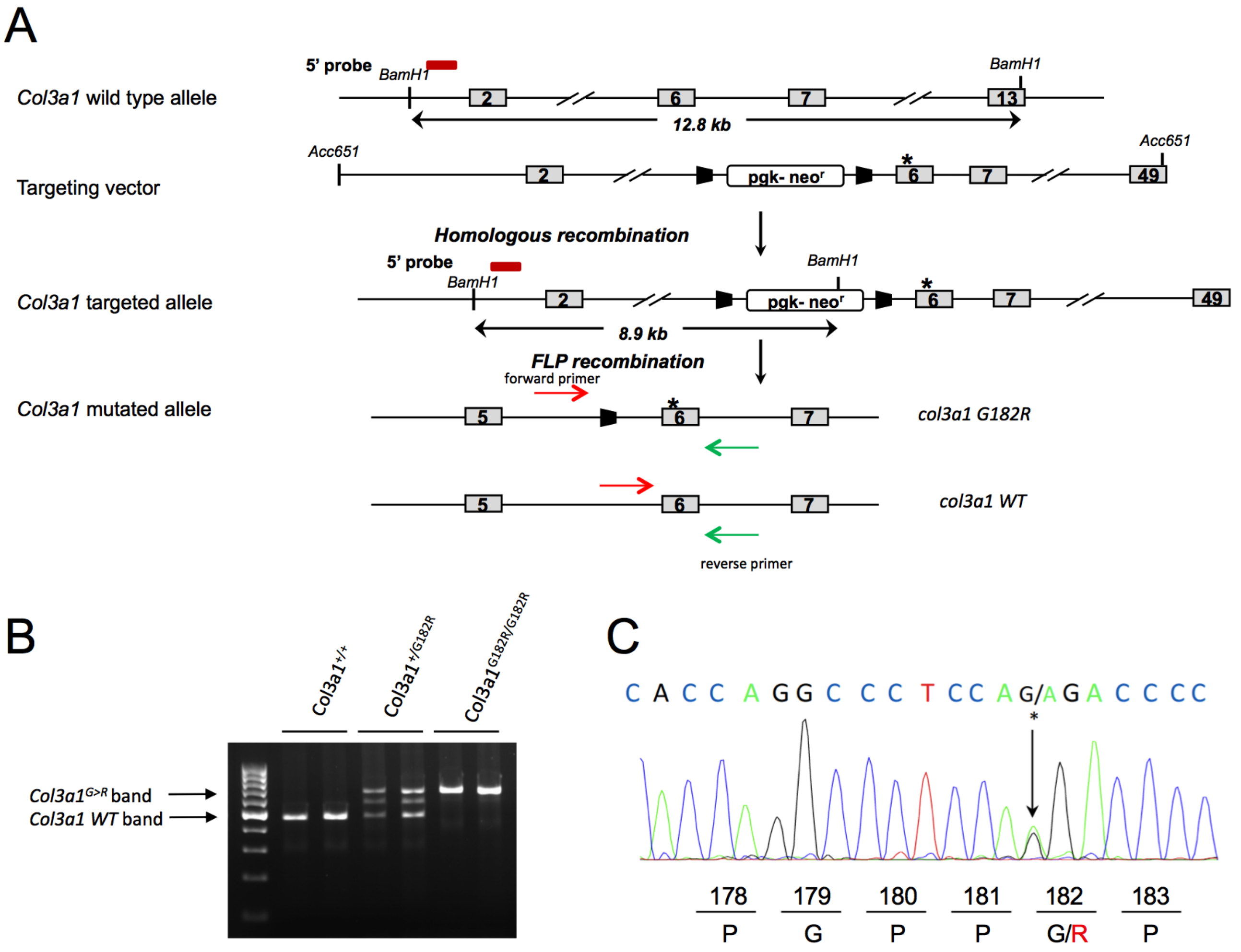


**Supplemental Figure 1: Gene targeting strategy and ES screen for *Col3a1* knock-in mice generation.**

**A.** Gene targeting strategy used for generating the *Col3a1*$^{+/G182R}$ mutated allele. The WT and targeted alleles are shown before and after FLP recombination (with and without the *pgk-neo*$^{r}$ cassette respectively). Boxes represent exons (the number of exon is indicated) and those with *pgk-neo*$^{r}$ represent the neomycin selection cassette. Black boxes correspond to FRT recombination sites, the asterisk represents the mutated site in exon 6. For Southern blot analysis, the 5' probe used is shown as a red rectangle upper the *Col3a1* WT and targeted allele and relevant restrictions sites are shown (*BamH1*). For genotype analysis, the *col3a1* reverse (green arrows) and forward (red arrows) primers are shown on both WT and mutated allele.

**B.** The G to A substitution is checked in ES cell clones by Sanger sequencing. Compared with WT ES cell clones, heterozygous ES cell clones reveal the substitution c.547G>A leading to an amino-acid change G182R. The asterisk represents the mutated nucleotide on exon 6.

**C.** Genotype analysis of WT and *Col3a1* mutated mice. Both WT and mutated alleles were detected by PCR using *Col3a1* primers.

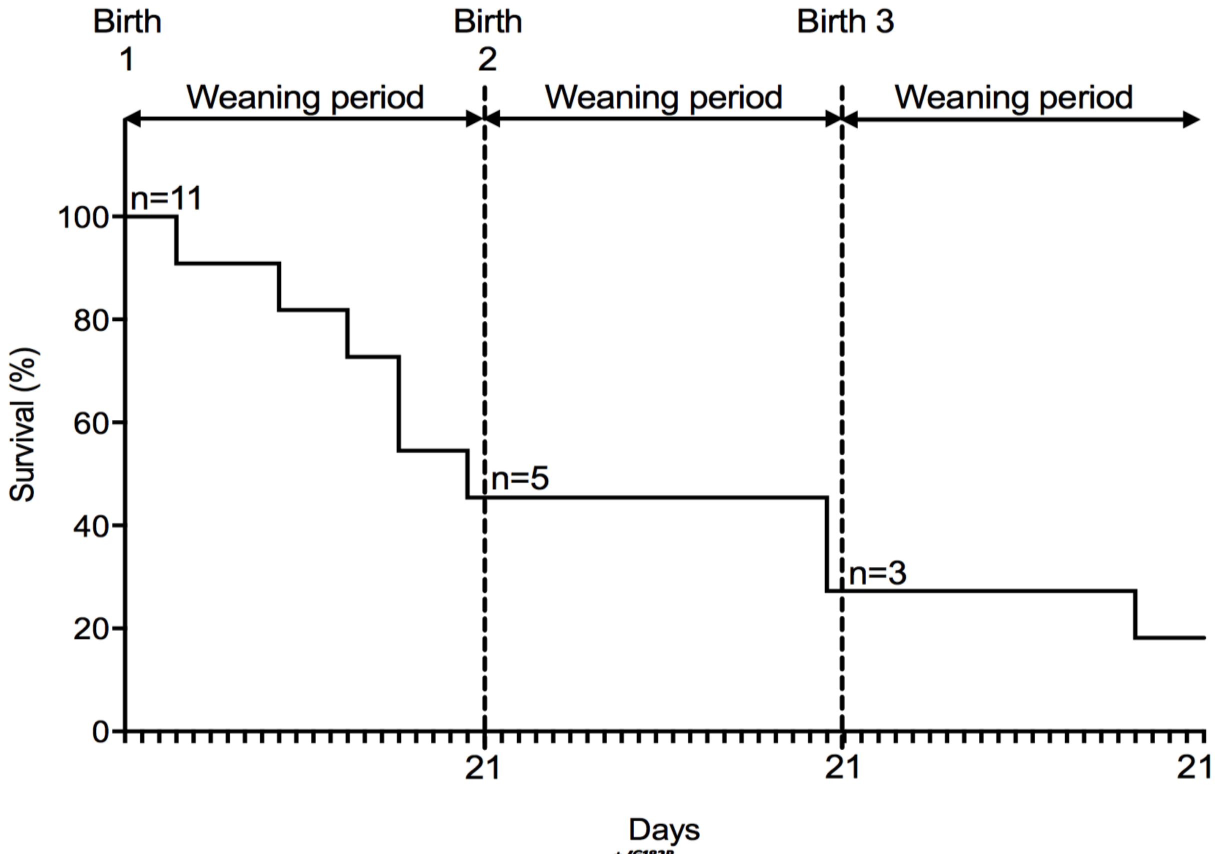


**Supplemental Figure 2 : Survival rate of *Col3a1*$^{+/G182R}$ female mice in weaning period.**

Kaplan-Meier Survival curve of *Col3a1*$^{+/G182R}$ female mice (n=11) in three successive weaning periods (21 days for each period). Six, two and one deaths are observed during the first, second and third weaning periods respectively. Thoracic aortic rupture is observed at autopsy for 90.9% (n=10/11) of mice.

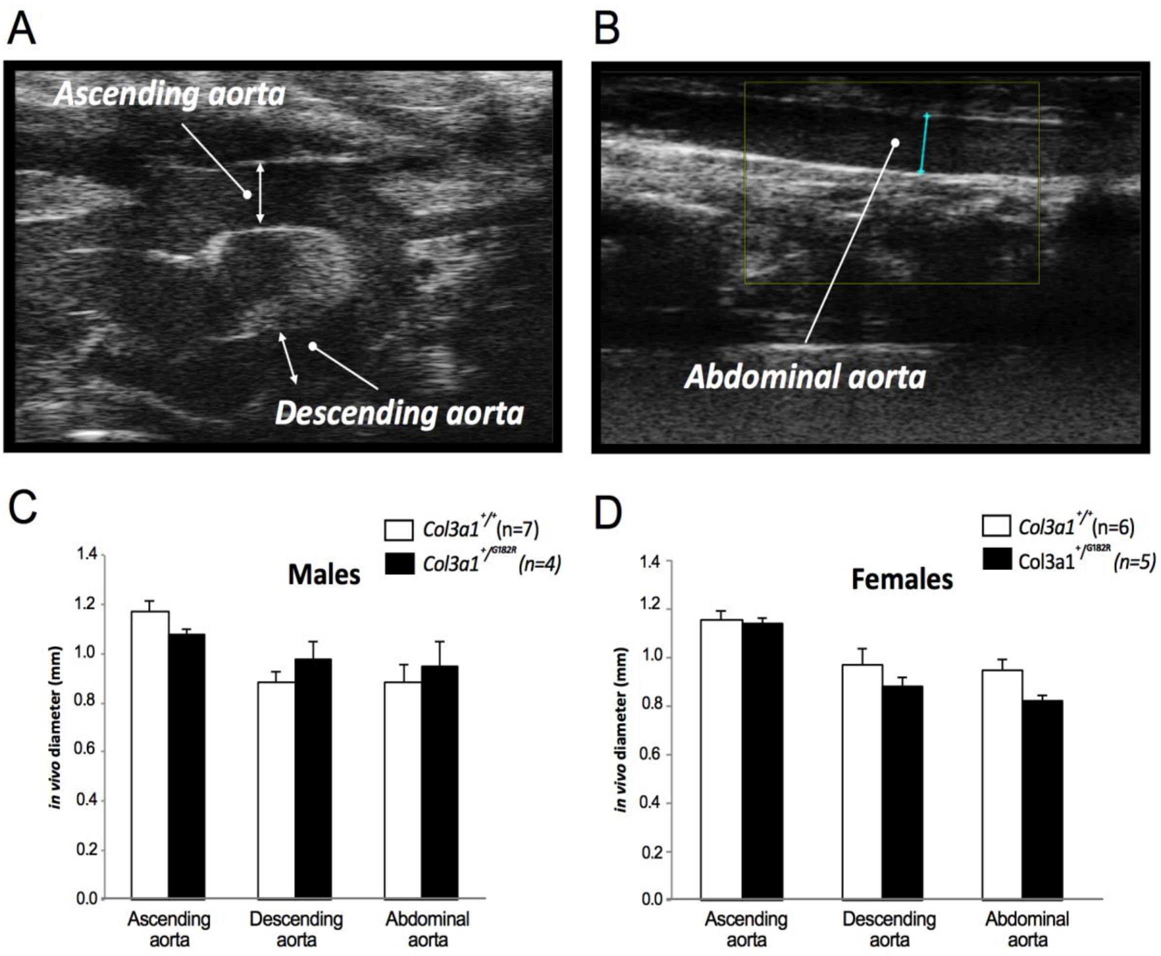


**Supplemental Figure 3:** ***in vivo*** **aorta diameter measured in 6-week-old** ***Col3a1***$^{+/+}$ **and** ***Col3a1***$^{+/G182R}$ **mice by echography.**

**A.** Ascending and descending thoracic aorta in *Col3a1*$^{+/G182R}$. White arrows indicate the ascending and descending thoracic aortic diameter.

**B.** Abdominal aorta in *Col3a1*$^{+/G182R}$. Blue arrow indicates the abdominal aortic diameter.

**C-D.** Measurement of the three diameters in males and females respectively. Error bars show mean ± SEM. No significant differences were found using Student t-test when the three diameters were compared between *Col3a1*$^{+/G182R}$ and *Col3a1*$^{+/+}$ mice in both sexes ($p>0.05$).

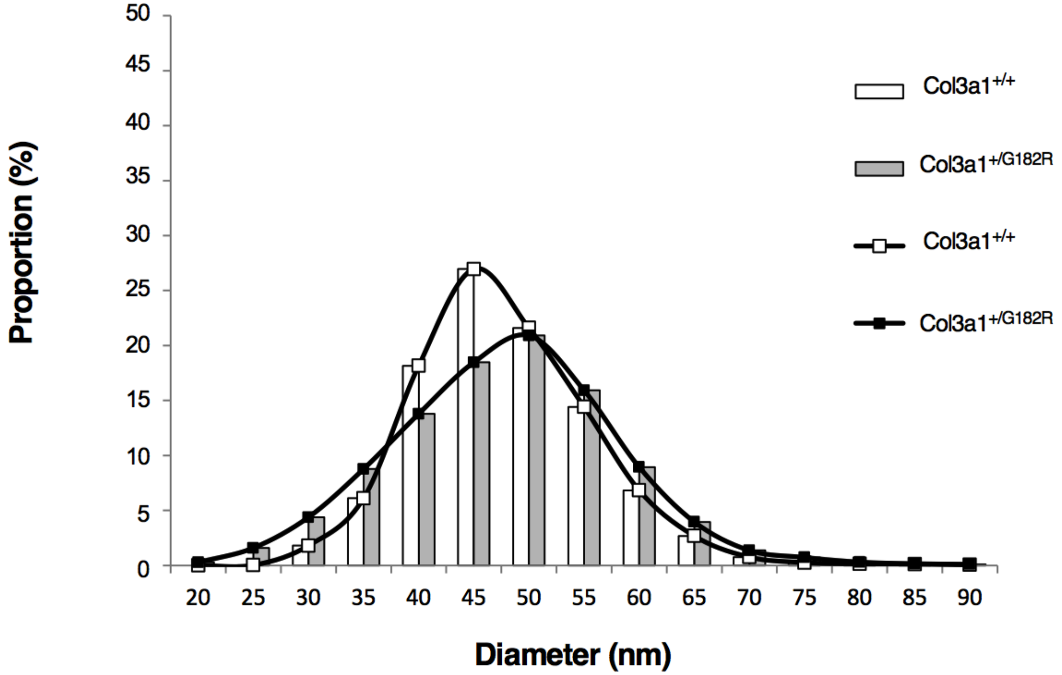


| | Col3a1$^{+/+}$ | Col3a1$^{+/G182R}$ |
|---|---|---|
| **min** | 24.29 | 9.24 |
| **max** | 101.97 | 118.92 |
| **mean** | 45.24 | 45.09 |
| **sd** | 8.10 | 10.27 |
| **nb fibres** | 4219 | 3611 |
| **% fibres** | 100 | 85.59 |

**Supplemental Figure 4: Heterogeneity of collagen fibers in *Col3a1*$^{+/G182R}$ mice.**

The proportion (%) of collagen fibrils with a given diameter (nm) shows a heterogeneity in the distribution for *Col3a1*$^{+/G182R}$ mice compared to *Col3a1*$^{+/+}$ controls in the aorta. In the table, the diameters of the collagen fibrils show a wider range in *Col3a1*$^{+/G182R}$ mice than in *Col3a1*$^{+/+}$ mice. The % fibers represent the number of collagen fibrils per unit area considering the number of collagen fibrils in *Col3a1*$^{+/+}$ mice as the reference. The % fibers reveal a lower density of collagen fibrils in *Col3a1*$^{+/G182R}$ mice than in *Col3a1*$^{+/+}$ mice.

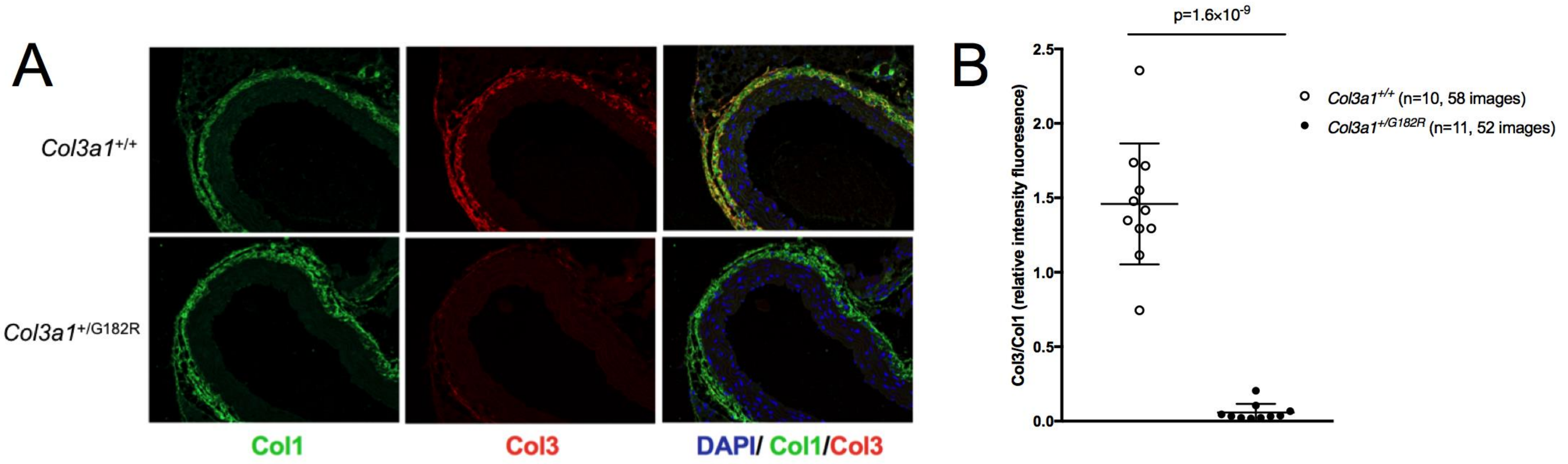


**Supplemental Figure 6 : Immunofluorescence of descending TA sections in *Col3a1*$^{+/G182R}$ mice.**

**A** Immunofluorescent staining of descending TA sections of *Col3a1*$^{+/+}$ (n=10) and *Col3a1*$^{+/G182R}$ (n=10) mice to show the distribution of collagens I and III. The sections are stained with collagen I (green) and collagen III (red) polyclonal antibodies and the cell nuclei were stained with DAPI (blue). The anti-collagen III antibody failed to recognize mature collagen III in *Col3a1*$^{+/G182R}$ mice.

**B** Relative quantification of the mature collagen III using the Collagen III/Collagen I immunofluorescence intensity ratio : the quantity of collagen III was collapsed probably due to conformational changes which lead to the absence of detection of collagen III, in the TA of *Col3a1*$^{+/G182R}$ mice compared to *Col3a1*$^{+/+}$ controls (Student t-test, $p=1.6\times10^{-9}$).

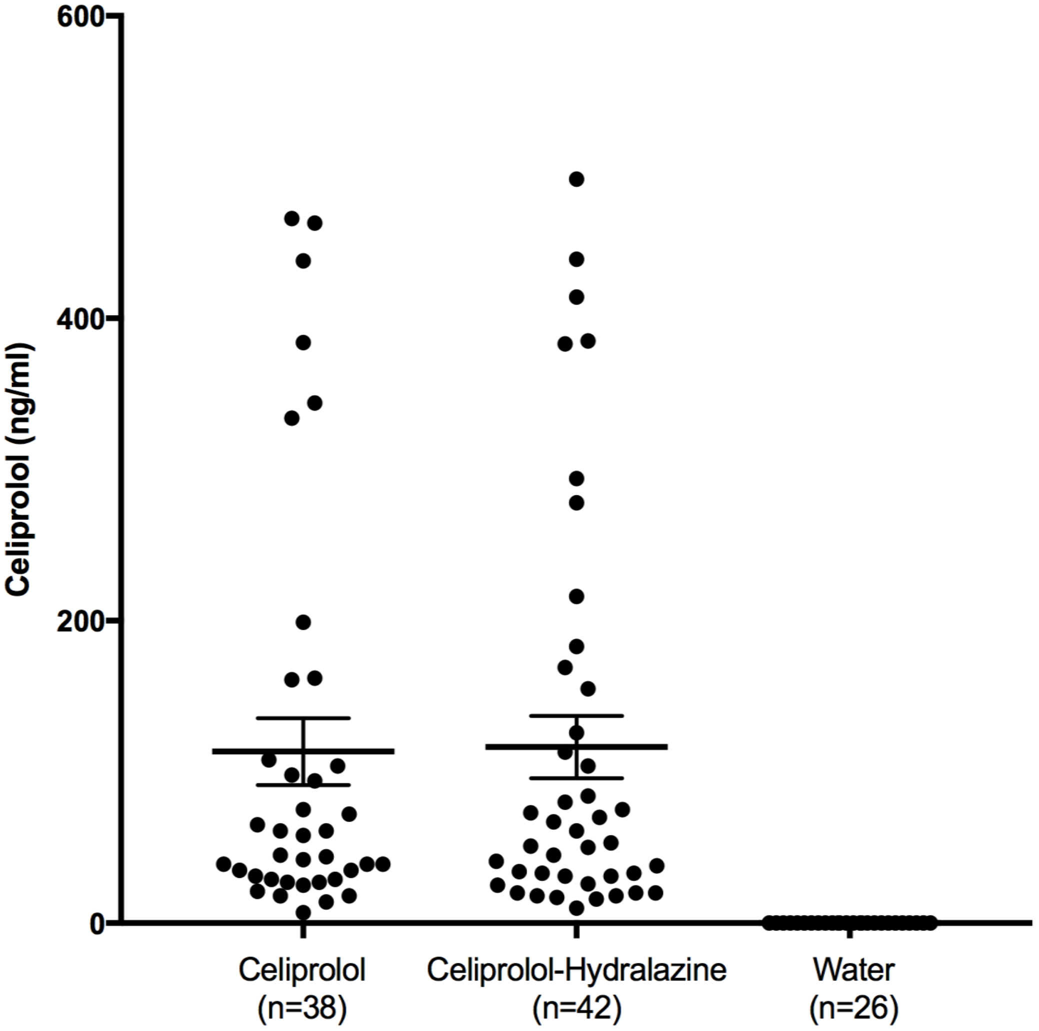


**Supplemental Figure 7 : Plasma celiprolol concentration measurement.**

Plasma concentration of celiprolol in *Col3a1*$^{+/G182R}$ (n=80) mice receiving celiprolol or the association celiprolol-hydralazine and in *Col3a1*$^{+/G182R}$ (n=26) mice receiving water, all of them being included in the celiprolol and celiprolol-hydralazine protocols. The celiprolol concentration was not significantly difference between the two treated groups.

Data are expressed as the mean ± SEM.

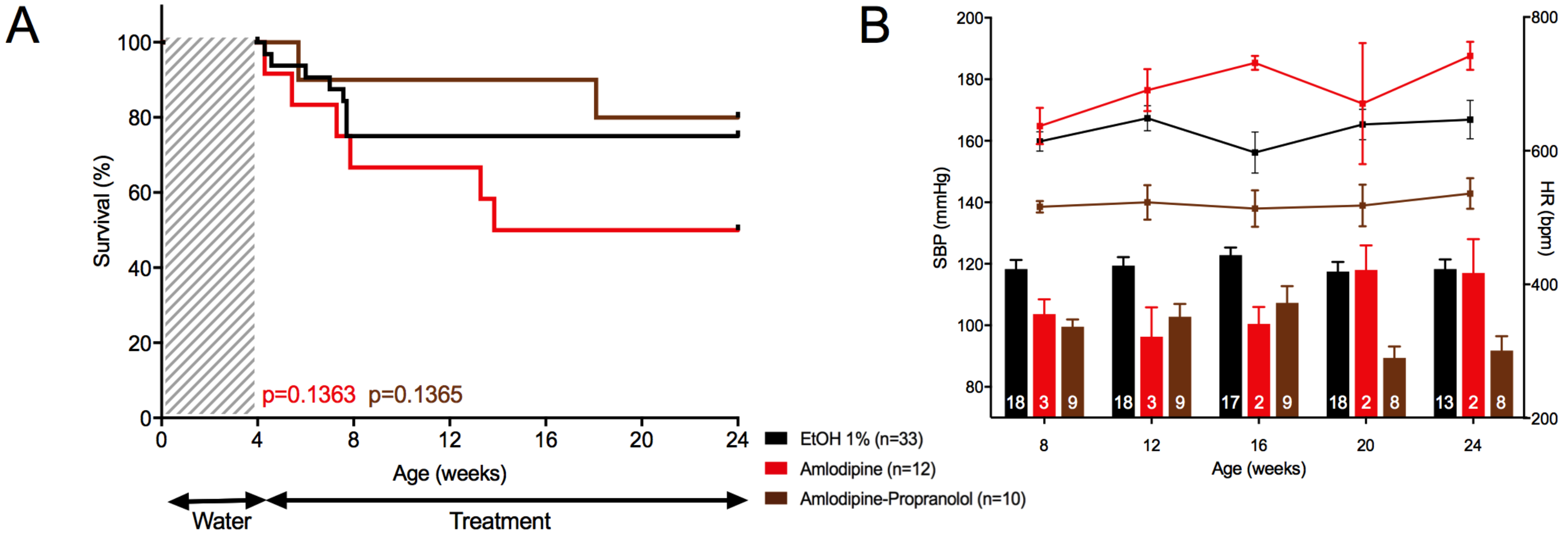


**Supplemental Figure 8 : Consequences of treatment by amlodipine and the association amlodipine – propranolol in *Col3a1*$^{+/G182R}$ female mice.**

**A** Survival comparison between Amlodipine and ethanol 1%, and between the association amlodipine-propranolol and amlodipine (monotherapy).

Kaplan-Meier Survival curve (red curve) for comparing *Col3a1* $^{+/G182R}$ treated with amlodipine (n=12) to *Col3a1* $^{+/G182R}$ treated with ethanol 1% (n=33). Amlodipine worsens the mortality despite no significant difference is observed using Log-Rank (Mantel-Cox) analysis (p=0.1363).

Kaplan-Meier Survival curve (brown curve) for comparing *Col3a1* $^{+/G182R}$ treated with amlodipine-propranolol (n=10) to *Col3a1* $^{+/G182R}$ treated with amlodipine (n=12). The association improves slightly the survival despite insignificant difference is calculated using Log-Rank (Mantel-Cox) analysis (p=0.1365).

**B** SBP and HR comparison between amlodipine and ethanol 1%, and between the association amlodipine-propranolol and amlodipine (monotherapy).

A significant decrease in SBP (linear mixed-effects model, $p=8.17\times10^{-3}$) was observed that was associated with no change in HR (student t-test, p>0.05 for each time of the 24-week follow-up period) when comparing amlodipine to ethanol 1% (red bars and curve).

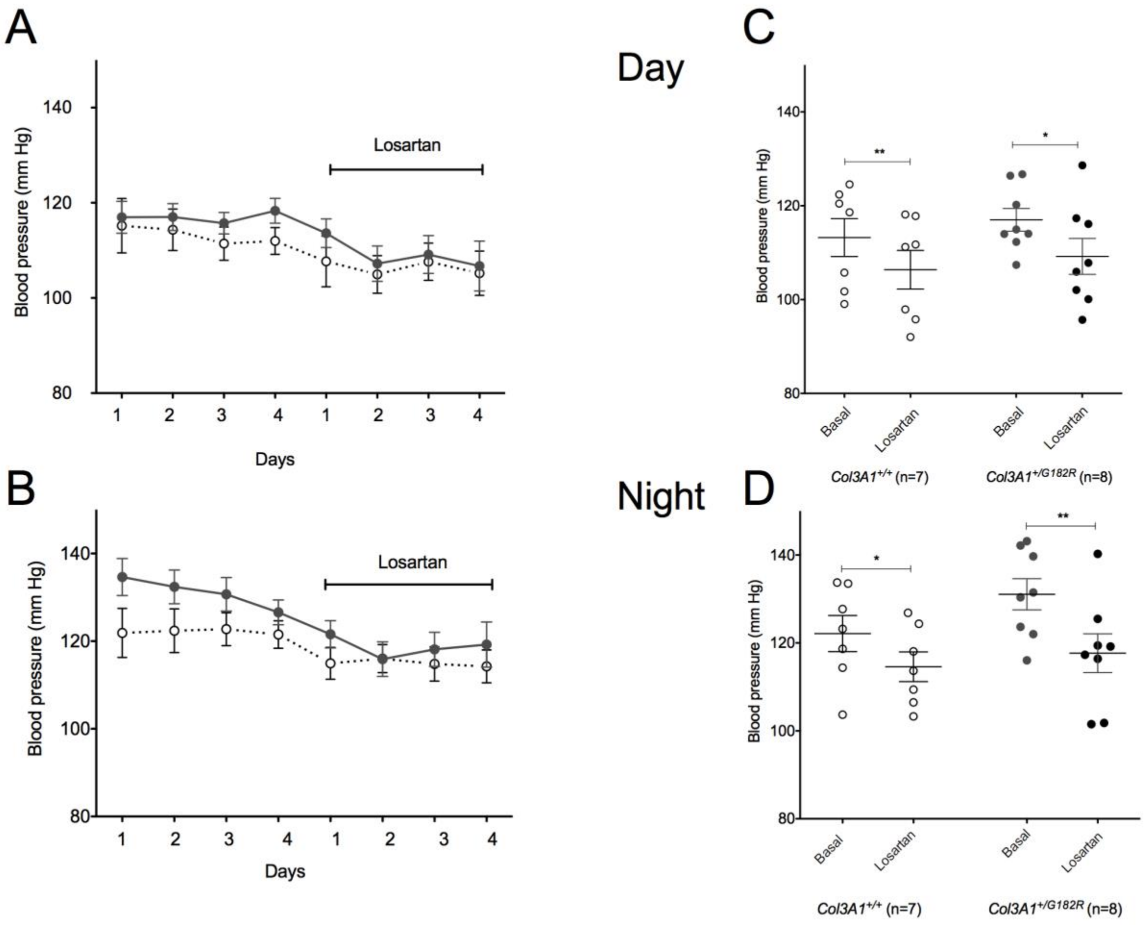


**Supplemental Figure 9: Normal BP in *Col3a1*$^{+/G182R}$ mice and decrease on Losartan.**

**A** Day SBP before (4 days= basal) or during oral administration (4 days) of losartan (135 mg/kg/day) in *Col3a1*$^{+/G182R}$ mice (n=8) compared to *Col3a1*$^{+/+}$ mice (n=7) examined with a telemetric system.

**B** Night SBP before (4 days= basal) or during oral administration (4 days) of losartan (135 mg/kg/day) in the same *Col3a1*$^{+/G182R}$ mice (n=8) and *Col3a1*$^{+/+}$ mice (n=7).

**C** Comparison of the mean of the day SBP before and during administration of losartan in the same *Col3a1*$^{+/G182R}$ mice (n=8) and *Col3a1*$^{+/+}$ mice (n=7). Significant decrease is observed in both groups.

**D** Comparison of the mean of the night SBP before and during administration of losartan in the same *Col3a1*$^{+/G182R}$ mice (n=8) and *Col3a1*$^{+/+}$ mice (n=7). Significant decrease is observed in both groups.

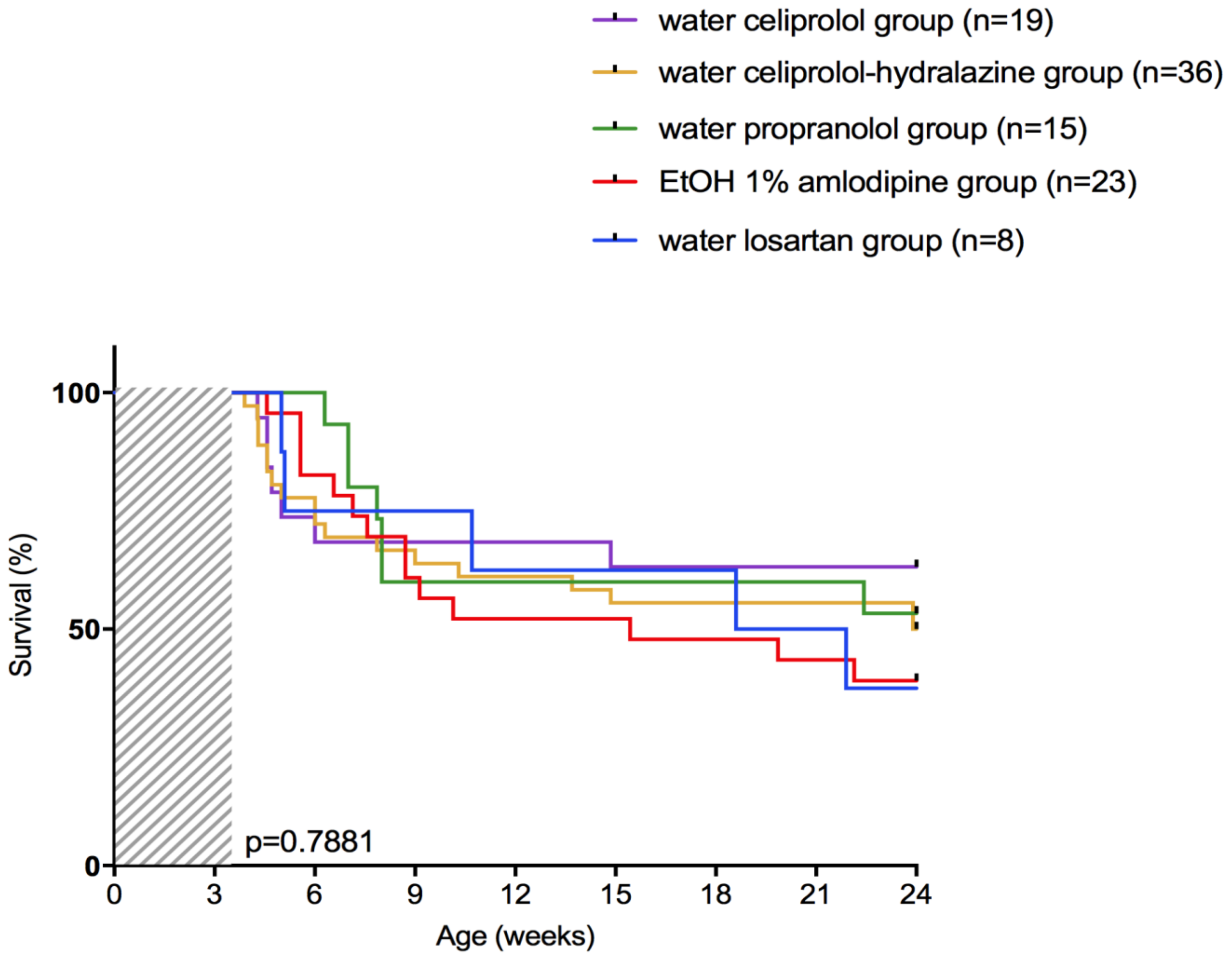


**Supplemental Figure 10 : Independent groups of *Col3a1*$^{+/G182R}$ control mice performed over time.**
Kaplan-Meier Survival curve for comparing the 5 *Col3a1*$^{+/G182R}$ male control groups. No significant difference is observed using Log-Rank (Mantel-Cox) analysis (p=0.7881).

**Supplemental Table 1: Primers used for the generation of the knock-in model, the genotyping of *Col3a1*$^{+/G182R}$ *and Col3a1*$^{+/+}$ mice, the droplet PCR and the RT-qPCR.**

| Name | Sequence 5'→ 3' |
|---|---|
| Construction | |
| SalI-ex4LA-fw | gtcgacgatatttccacaatatgcttatcca |
| NdeI/BsrgI-in4LA-rv | catatgtgtacaatttttgcatgaaaactttactt |
| NotI-in4RA-fw | gcggccgcctttattgaaaatgtcccagaa |
| SalI-in5RA-rv | gtcgacgtggatgtaggcagaaatttta |
| ex5RA-fw | cttcatgaatatatgaagcaagc |
| SalI-in5RA-rv | gtcgacgtggatgtaggcagaaatttta |
| Ex6-point-mutation-fw | ccaggccctccaagaccccctggtt |
| Ex6-point-mutation-rv | aaccagggggtcttggagggcctgg |
| XbaI-LA-gap-repair-fw | tctagaTGCCAATGAGAATTGACCAT |
| BglII-LA-gap-repair-rv | agatctTTGGAATGTTTACGCTGACTC |
| BglII-RA-gap-repair-fw | agatctCAAAGATGGATCACCTGGAG |
| NcoI-RA-gap-repair-rv | ccatggCTCTAAATTTGGGGCCAGTC |
| Genotyping | |
| Col3a1KI-fw | tcatctgaagtaaagttttcatgc |
| Col3a1KI-rv | tttcaccgaaattgagtggtt |
| Droplet Digital PCR | |
| Col3a1KI-ddpcr-fw | TGTCAAGTCTGGAGTGGGAG |
| Col3a1KI-ddpcr-rv | CAGGATGTCCAGAAGAACCA |
| Col3a1wt-probe | ACCAGGCCCTCCA**G**GACCC |
| Col3a1KI-probe | ACCAGGCCCTCCA**A**GACCC |
| Vimentin | Bio-Rad cat#dMmuCPE5097200 |
| qPCR | |
| Atf6-fw | TGCCTTGGGAGTCAGACCTAT |
| Atf6-rv | GCTGAGTTGAAGAACACGAGTC |
| ATG5-fw | TCCTCGCTAGATGGAACCAC |
| ATG5-rv | AGTGGTCCTGTGTGTCTCAG |
| Atg7-fw | CCAGTCCGTTGAAGTCCTCT |
| Atg7-rv | AGATGACTCAGCCAGCCTTT |
| Hsp47-fw | ACCCCTTCATCTTCCTGGTG |
| Hsp47-rv | CCCATGTGTCTCAGGAACCT |

**Supplemental Table 2 : Basic parameters (weight, SBP and HR) in *Col3a1*$^{+/G182R}$ and *Col3a1*$^{+/+}$ mice throughout the 24-week follow-up period.**

| | | Males | | | Females | | | All | | |
|---|---|---|---|---|---|---|---|---|---|---|
| | Age (weeks) | col3a1$^{+/+}$ | col3a1$^{+/G182R}$ | p value | col3a1$^{+/+}$ | col3a1$^{+/G182R}$ | p value | col3a1$^{+/+}$ | col3a1$^{+/G182R}$ | p value |
| Weight (g) | 5 | 18.2 ± 0.4<br>n = 29 | 16.4 ± 0.4<br>n = 36 | 0.002 | 14.8 ± 0.3<br>n = 25 | 14.8 ± 0.3<br>n = 32 | 0.934 | 16.6 ± 0.4<br>n = 54 | 15.6 ± 0.2<br>n = 68 | 0.020 |
| | 8 | 23.0 ± 0.4<br>n = 36 | 21.5 ± 0.4<br>n = 29 | 0.008 | 17.6 ± 0.3<br>n = 24 | 17.3 ± 0.2<br>n = 32 | 0.274 | 20.9 ± 0.4<br>n = 60 | 19.3 ± 0.3<br>n = 61 | 0.005 |
| | 12 | 26.3 ± 0.4<br>n = 36 | 25.1 ± 0.4<br>n = 22 | 0.071 | 20.3 ± 0.3<br>n = 26 | 19.6 ± 0.2<br>n = 32 | 0.043 | 23.8 ± 0.4<br>n = 62 | 21.8 ± 0.4<br>n = 54 | 0.003 |
| | 16 | 28.3 ± 0.5<br>n = 36 | 26.7 ± 0.5<br>n = 21 | 0.020 | 22.1 ± 0.3<br>n = 26 | 21.3 ± 0.3<br>n = 32 | 0.052 | 25.7 ± 0.5<br>n = 62 | 23.4 ± 0.4<br>n = 53 | 0.001 |
| | 20 | 29.3 ± 0.4<br>n = 35 | 27.9 ± 0.5<br>n = 21 | 0.045 | 22.8 ± 0.3<br>n = 26 | 22.1 ± 0.3<br>n = 31 | 0.109 | 26.5 ± 0.5<br>n = 61 | 24.4 ± 0.5<br>n = 52 | 0.003 |
| | 24 | 30.3 ± 0.5<br>n = 35 | 28.6 ± 0.6<br>n = 18 | 0.062 | 23.6 ± 0.3<br>n = 26 | 22.6 ± 0.4<br>n = 30 | 0.064 | 27.4 ± 0.5<br>n = 61 | 24.9 ± 0.5<br>n = 48 | 0.001 |
| SBP (mmHg) | 8 | 119.7 ± 2.7<br>n = 37 | 116.5 ± 3.1<br>n = 27 | 0.431 | 121.2 ± 3.4<br>n = 26 | 117.0 ± 2.2<br>n = 32 | 0.284 | 120.3 ± 2.1<br>n = 63 | 116.8 ± 1.8<br>n = 59 | 0.200 |
| | 12 | 113.6 ± 2.2<br>n = 37 | 115.3 ± 2.6<br>n = 20 | 0.647 | 118.5 ± 3.4<br>n = 24 | 111.8 ± 2.4<br>n = 29 | 0.102 | 115.5 ± 1.9<br>n = 61 | 113.2 ± 1.8<br>n = 49 | 0.415 |
| | 16 | 119.3 ± 2.2<br>n = 36 | 117.93 ± 2.4<br>n = 20 | 0.680 | 118.0 ± 3.2<br>n = 24 | 112.7 ± 2.4<br>n = 28 | 0.182 | 118.8 ± 1.8<br>n = 60 | 114.9 ± 1.8<br>n = 48 | 0.126 |
| | 20 | 115.1 ± 2.1<br>n = 34 | 111.8 ± 3.2<br>n = 20 | 0.368 | 112.5 ± 2.4<br>n = 25 | 113.3 ± 2.3<br>n = 31 | 0.802 | 114.0 ± 1.6<br>n = 59 | 112.7 ± 1.9<br>n = 51 | 0.571 |
| | 24 | 120.7 ± 3.0<br>n = 26 | 112.2 ± 3.4<br>n = 13 | 0.094 | 115.1 ± 3.0<br>n = 22 | 108.9 ± 2.8<br>n = 21 | 0.135 | 118.2 ± 2.2<br>n = 48 | 110.2 ± 2.1<br>n = 34 | 0.012 |
| HR (bpm) | 8 | 653.7 ± 10.9<br>n = 37 | 625.4 ± 625.4<br>n = 27 | 0.115 | 607.2 ± 17.2<br>n = 26 | 585.3 ± 13.7<br>n = 32 | 0.319 | 634.6 ± 9.9<br>n = 63 | 603.7 ± 10.2<br>n = 59 | 0.032 |
| | 12 | 656.5 ± 10.2<br>n = 37 | 636.6 ± 13.1<br>n = 20 | 0.242 | 609.4 ± 15.3<br>n = 24 | 610.5 ± 15.4<br>n = 29 | 0.961 | 638.0 ± 9.0<br>n = 61 | 621.2 ± 10.7<br>n = 49 | 0.228 |
| | 16 | 657.1 ± 10.5<br>n = 36 | 638.1 ± 13.4<br>n = 20 | 0.280 | 594.6 ± 17.5<br>n = 24 | 605.9 ± 14.6<br>n = 28 | 0.621 | 632.1 ± 10.2<br>n = 60 | 619.3 ± 10.3<br>n = 48 | 0.385 |
| | 20 | 684.2 ± 8.6<br>n = 34 | 659.8 ± 12.4<br>n = 20 | 0.103 | 601.0 ± 18.7<br>n = 25 | 567.8 ± 17.5<br>n = 31 | 0.201 | 649.0 ± 10.7<br>n = 59 | 603.8 ± 13.2<br>n = 51 | 0.009 |
| | 24 | 667.4 ± 10.5<br>n = 26 | 673.0 ± 13.4<br>n = 13 | 0.750 | 646.5 ± 14.3<br>n = 22 | 607.9 ± 15.3<br>n = 21 | 0.072 | 657.8 ± 8.7<br>n = 48 | 632.8 ± 12.0<br>n = 34 | 0.086 |

Data are expressed as the mean ± SEM. The number of mice at each time are indicated above the mean ± SEM.
Student-t test.

**Supplemental Table 3 : Differential expression of the 223 genes belonging to the MAPK signaling pathway of KEGG database (corresponding to the PLC/IP3/PKC/ERK signaling pathway) in the transcriptomic data of *Col3a1*$^{+/G182R}$ mice.**

| Gene symbol | Mean Col3a1$^{+/G182R}$ (n=3) | Mean Col3a1$^{+/+}$ (n=4) | log2FoldChange | FoldChange | p-value | *adjusted p-value |
|---|---|---|---|---|---|---|
| AKT1 | 3317.42 | 2918.34 | 0.18 | 1.14 | 0.313 | 0.561 |
| AKT2 | 3747.25 | 3344.54 | 0.16 | 1.12 | 0.220 | 0.472 |
| AKT3 | 2631.11 | 1878.04 | 0.49 | 1.40 | 0.091 | 0.317 |
| ARRB1 | 1045.41 | 956.75 | 0.13 | 1.09 | 0.559 | 0.756 |
| ARRB2 | 233.26 | 299.98 | -0.36 | 0.78 | 0.239 | 0.492 |
| ATF2 | 1101.47 | 875.86 | 0.33 | 1.26 | 0.021 | 0.176 |
| ATF4 | 5051.78 | 5289.14 | -0.07 | 0.96 | 0.607 | 0.791 |
| BDNF | 277.01 | 114.65 | 1.27 | 2.41 | 0.003 | 0.083 |
| BRAF | 1415.40 | 1103.65 | 0.36 | 1.28 | 0.137 | 0.378 |
| CACNA1A | 45.37 | 48.61 | -0.11 | 0.92 | 0.799 | 0.902 |
| CACNA1B | | | | | | |
| CACNA1C | 2255.52 | 2386.02 | -0.08 | 0.95 | 0.723 | 0.861 |
| CACNA1D | 357.95 | 466.54 | -0.38 | 0.77 | 0.205 | 0.456 |
| CACNA1E | 28.36 | 44.11 | -0.57 | 0.68 | 0.259 | 0.511 |
| CACNA1F | | | | | | |
| CACNA1G | 135.82 | 128.80 | 0.07 | 1.05 | 0.905 | 0.956 |
| CACNA1H | 42.04 | 36.24 | 0.14 | 1.10 | 0.755 | 0.879 |
| CACNA1I | | | | | | |
| CACNA1S | | | | | | |
| CACNA2D1 | 2364.49 | 1892.23 | 0.32 | 1.25 | 0.216 | 0.467 |
| CACNA2D2 | 4.50 | 5.28 | -0.12 | 0.92 | 0.902 | NA |
| CACNA2D3 | 46.73 | 27.88 | 0.72 | 1.65 | 0.113 | 0.349 |
| CACNA2D4 | 16.34 | 12.51 | 0.46 | 1.37 | 0.496 | NA |
| CACNB1 | 189.48 | 267.25 | -0.49 | 0.71 | 0.036 | 0.217 |
| CACNB2 | 767.78 | 697.66 | 0.14 | 1.10 | 0.440 | 0.667 |
| CACNB3 | 912.75 | 1164.61 | -0.35 | 0.78 | 0.084 | 0.308 |
| CACNB4 | 47.51 | 40.30 | 0.23 | 1.17 | 0.462 | 0.684 |
| CACNG1 | | | | | | |
| CACNG2 | | | | | | |
| CACNG3 | | | | | | |
| CACNG4 | 39.36 | 37.94 | 0.03 | 1.02 | 0.953 | 0.978 |
| CACNG5 | | | | | | |
| CACNG6 | | | | | | |
| CACNG7 | 468.00 | 397.58 | 0.23 | 1.18 | 0.426 | 0.655 |
| CACNG8 | | | | | | |
| CASP3 | 505.37 | 440.88 | 0.20 | 1.15 | 0.277 | 0.530 |
| CD14 | 107.24 | 88.85 | 0.27 | 1.21 | 0.325 | 0.572 |
| CDC25B | 371.68 | 562.33 | -0.60 | 0.66 | 0.061 | 0.267 |
| CDC42 | 8374.08 | 7340.68 | 0.19 | 1.14 | 0.294 | 0.544 |
| CHP1 | 3792.95 | 2861.99 | 0.41 | 1.33 | 0.130 | 0.372 |
| CHP2 | 89.49 | 46.77 | 0.93 | 1.91 | 0.060 | 0.265 |
| CHUK | 714.51 | 920.96 | -0.37 | 0.78 | 0.089 | 0.314 |
| CRK | 2975.77 | 2025.29 | 0.55 | 1.47 | 0.061 | 0.268 |
| CRKL | 1138.22 | 753.71 | 0.59 | 1.51 | 0.015 | 0.157 |
| DAXX | 369.85 | 507.32 | -0.45 | 0.73 | 0.019 | 0.167 |
| DDIT3 | 549.77 | 773.30 | -0.49 | 0.71 | 0.015 | 0.156 |
| DUSP1 | 2921.22 | 1900.58 | 0.62 | 1.54 | 0.036 | 0.217 |
| **DUSP10** | **498.74** | **320.20** | **0.64** | **1.56** | **1.498E-04** | **0.025** |
| DUSP14 | 184.74 | 188.70 | -0.03 | 0.98 | 0.889 | 0.947 |
| DUSP16 | 700.22 | 517.22 | 0.44 | 1.36 | 0.014 | 0.149 |
| DUSP2 | 23.28 | 7.34 | 1.67 | 3.18 | 0.024 | NA |
| DUSP3 | 4610.89 | 3700.79 | 0.32 | 1.25 | 0.317 | 0.565 |
| DUSP4 | 61.06 | 31.01 | 0.99 | 1.98 | 0.124 | 0.363 |
| DUSP5 | 77.08 | 71.03 | 0.12 | 1.09 | 0.714 | 0.855 |
| DUSP6 | 383.76 | 242.84 | 0.66 | 1.58 | 0.129 | 0.370 |
| DUSP7 | 400.79 | 514.50 | -0.36 | 0.78 | 0.031 | 0.206 |
| DUSP8 | 1187.52 | 741.59 | 0.68 | 1.60 | 0.004 | 0.091 |
| DUSP9 | | | | | | |
| ECSIT | 469.11 | 563.55 | -0.26 | 0.83 | 0.159 | 0.405 |
| EGF | 16.74 | 35.21 | -1.05 | 0.48 | 0.011 | 0.136 |
| EGFR | 1042.45 | 723.47 | 0.53 | 1.44 | 0.001 | 0.060 |

| Gene symbol | Mean Col3a1$^{+/G182R}$ (n=3) | Mean Col3a1$^{+/+}$ (n=4) | log2FoldChange | FoldChange | p-value | *adjusted p-value |
|---|---|---|---|---|---|---|
| ELK1 | 428.76 | 355.90 | 0.27 | 1.20 | 0.259 | 0.512 |
| ELK4 | 887.17 | 1279.10 | -0.53 | 0.69 | 0.007 | 0.114 |
| FAS | 2029.35 | 2008.94 | 0.01 | 1.01 | 0.949 | 0.976 |
| FASLG | | | | | | |
| FGF1 | 2813.85 | 2510.74 | 0.16 | 1.12 | 0.328 | 0.574 |
| FGF10 | 9.97 | 5.20 | 1.09 | 2.13 | 0.141 | NA |
| FGF11 | 297.97 | 363.61 | -0.29 | 0.82 | 0.086 | 0.310 |
| FGF12 | 3.79 | 10.93 | -1.60 | 0.33 | 0.160 | NA |
| FGF13 | 281.96 | 184.16 | 0.61 | 1.53 | 0.106 | 0.340 |
| FGF14 | 573.31 | 512.99 | 0.16 | 1.12 | 0.596 | 0.784 |
| FGF16 | 65.59 | 63.08 | 0.04 | 1.03 | 0.900 | 0.953 |
| FGF17 | 1.81 | 5.28 | -1.59 | 0.33 | 0.098 | NA |
| FGF18 | 15.76 | 15.94 | -0.03 | 0.98 | 0.966 | 0.984 |
| FGF19 | | | | | | |
| FGF2 | 2475.66 | 1870.52 | 0.40 | 1.32 | 0.195 | 0.444 |
| FGF20 | | | | | | |
| FGF21 | | | | | | |
| FGF22 | | | | | | |
| FGF23 | | | | | | |
| FGF3 | | | | | | |
| FGF4 | | | | | | |
| FGF5 | | | | | | |
| FGF6 | | | | | | |
| FGF7 | 70.44 | 45.15 | 0.61 | 1.53 | 0.080 | 0.302 |
| FGF8 | | | | | | |
| FGF9 | 10.68 | 13.52 | -0.29 | 0.82 | 0.676 | NA |
| FGFR1 | 3440.86 | 3870.17 | -0.17 | 0.89 | 0.299 | 0.549 |
| FGFR2 | 2292.69 | 2656.85 | -0.21 | 0.86 | 0.410 | 0.643 |
| FGFR3 | 127.60 | 135.22 | -0.09 | 0.94 | 0.789 | 0.897 |
| FGFR4 | | | | | | |
| FLNA | 136128.01 | 171394.09 | -0.33 | 0.79 | 0.033 | 0.212 |
| FLNB | 3288.45 | 3862.59 | -0.23 | 0.85 | 0.099 | 0.331 |
| FLNC | 2854.93 | 2839.21 | 0.01 | 1.01 | 0.975 | 0.989 |
| FOS | 674.87 | 143.46 | 2.23 | 4.70 | 0.003 | 0.089 |
| GADD45A | 172.91 | 176.84 | -0.03 | 0.98 | 0.911 | 0.959 |
| GADD45B | 523.14 | 424.04 | 0.30 | 1.23 | 0.262 | 0.515 |
| GADD45G | 709.33 | 592.83 | 0.26 | 1.20 | 0.343 | 0.586 |
| GNA12 | 2140.67 | 1763.89 | 0.28 | 1.21 | 0.192 | 0.441 |
| GNG12 | 5518.93 | 3583.40 | 0.62 | 1.54 | 0.088 | 0.312 |
| GRB2 | 1452.39 | 1204.83 | 0.27 | 1.20 | 0.155 | 0.400 |
| HRAS | 852.99 | 1163.85 | -0.45 | 0.73 | 0.028 | 0.199 |
| **HSPA1A** | **2117.43** | **776.63** | **1.45** | **2.73** | **1.048E-04** | **0.021** |
| **HSPA1B** | **1746.02** | **438.12** | **1.99** | **3.98** | **3.448E-06** | **0.005** |
| HSPA1L | 372.51 | 311.65 | 0.26 | 1.20 | 0.170 | 0.417 |
| HSPA2 | 895.86 | 863.59 | 0.05 | 1.04 | 0.811 | 0.910 |
| HSPA6 | | | | | | |
| HSPA8 | 38567.40 | 30348.34 | 0.35 | 1.27 | 0.287 | 0.538 |
| HSPB1 | 4384.41 | 3842.58 | 0.19 | 1.14 | 0.487 | 0.704 |
| IKBKB | 1071.06 | 1211.10 | -0.18 | 0.89 | 0.373 | 0.612 |
| IKBKG | 951.89 | 741.59 | 0.36 | 1.28 | 0.043 | 0.232 |
| IL1A | | | | | | |
| IL1B | 6.01 | 2.86 | 0.98 | 1.97 | 0.398 | NA |
| IL1R1 | 1354.05 | 1287.44 | 0.07 | 1.05 | 0.678 | 0.834 |
| IL1R2 | | | | | | |
| JMJD7-PLA2G4B | | | | | | |
| JUN | 2036.19 | 1392.21 | 0.55 | 1.46 | 0.019 | 0.169 |
| JUND | 2202.83 | 1941.46 | 0.18 | 1.13 | 0.426 | 0.655 |
| KRAS | 1208.79 | 1008.72 | 0.26 | 1.20 | 0.265 | 0.517 |
| LAMTOR3 | 1216.83 | 1092.77 | 0.15 | 1.11 | 0.225 | 0.478 |
| MAP2K1 | 1120.25 | 994.45 | 0.17 | 1.13 | 0.420 | 0.650 |
| MAP2K2 | 1232.57 | 1521.54 | -0.30 | 0.81 | 0.141 | 0.384 |

| Gene symbol | Mean Col3a1$^{+/G182R}$ (n=3) | Mean Col3a1$^{+/+}$ (n=4) | log2FoldChange | FoldChange | p-value | *adjusted p-value |
|---|---|---|---|---|---|---|
| **MAP2K3** | **1376.59** | **1738.40** | **-0.34** | **0.79** | **3.216E-05** | **0.012** |
| MAP2K4 | 1146.74 | 982.49 | 0.22 | 1.17 | 0.276 | 0.529 |
| MAP2K5 | 324.50 | 344.57 | -0.09 | 0.94 | 0.500 | 0.712 |
| MAP2K6 | 198.94 | 132.03 | 0.59 | 1.51 | 0.031 | 0.207 |
| MAP2K7 | 768.12 | 1008.92 | -0.39 | 0.76 | 0.125 | 0.365 |
| MAP3K1 | 762.69 | 733.92 | 0.05 | 1.04 | 0.835 | 0.921 |
| MAP3K11 | 376.29 | 348.83 | 0.11 | 1.08 | 0.658 | 0.822 |
| MAP3K12 | 564.52 | 577.39 | -0.03 | 0.98 | 0.830 | 0.919 |
| MAP3K13 | 74.53 | 65.47 | 0.19 | 1.14 | 0.422 | 0.652 |
| MAP3K14 | 117.53 | 140.14 | -0.25 | 0.84 | 0.275 | 0.527 |
| MAP3K2 | 1366.32 | 1139.14 | 0.26 | 1.20 | 0.106 | 0.340 |
| MAP3K20 | 4641.95 | 4558.34 | 0.03 | 1.02 | 0.832 | 0.920 |
| MAP3K3 | 1280.84 | 988.25 | 0.37 | 1.30 | 0.173 | 0.421 |
| MAP3K4 | 613.39 | 699.44 | -0.19 | 0.88 | 0.170 | 0.418 |
| MAP3K5 | 1063.23 | 1110.70 | -0.06 | 0.96 | 0.774 | 0.888 |
| MAP3K6 | 200.28 | 236.61 | -0.24 | 0.85 | 0.446 | 0.671 |
| MAP3K7 | 1521.03 | 1471.64 | 0.05 | 1.03 | 0.683 | 0.836 |
| MAP3K8 | 243.18 | 330.20 | -0.44 | 0.74 | 0.023 | 0.180 |
| MAP4K1 | 12.91 | 13.34 | -0.12 | 0.92 | 0.882 | 0.944 |
| MAP4K2 | 771.71 | 1368.85 | -0.83 | 0.56 | 0.005 | 0.098 |
| MAP4K3 | 726.31 | 674.21 | 0.11 | 1.08 | 0.488 | 0.704 |
| MAP4K4 | 2827.81 | 2948.69 | -0.06 | 0.96 | 0.697 | 0.845 |
| MAPK1 | 3174.48 | 2378.37 | 0.42 | 1.33 | 0.232 | 0.485 |
| MAPK10 | | | | | | |
| MAPK11 | 60.01 | 73.26 | -0.28 | 0.82 | 0.531 | 0.736 |
| MAPK12 | 333.67 | 592.47 | -0.83 | 0.56 | 0.015 | 0.154 |
| MAPK13 | 5.82 | 6.42 | -0.28 | 0.82 | 0.723 | NA |
| MAPK14 | 1791.45 | 2057.68 | -0.20 | 0.87 | 0.052 | 0.249 |
| MAPK3 | 2051.16 | 2105.50 | -0.04 | 0.97 | 0.757 | 0.880 |
| MAPK7 | 365.89 | 520.73 | -0.51 | 0.70 | 0.005 | 0.105 |
| MAPK8 | 688.91 | 488.57 | 0.49 | 1.41 | 0.052 | 0.249 |
| MAPK8IP1 | 431.79 | 401.46 | 0.10 | 1.07 | 0.503 | 0.716 |
| MAPK8IP2 | 5.86 | 2.74 | 1.01 | 2.02 | 0.472 | NA |
| MAPK8IP3 | 1835.18 | 2931.86 | -0.68 | 0.63 | 0.021 | 0.176 |
| MAPK9 | 1522.41 | 1240.02 | 0.30 | 1.23 | 0.111 | 0.347 |
| MAPKAPK2 | 2151.63 | 2010.06 | 0.10 | 1.07 | 0.401 | 0.636 |
| MAPKAPK3 | 144.82 | 157.74 | -0.12 | 0.92 | 0.731 | 0.866 |
| MAPKAPK5 | 701.05 | 793.76 | -0.18 | 0.88 | 0.249 | 0.503 |
| MAPT | 145.78 | 187.79 | -0.37 | 0.77 | 0.307 | 0.557 |
| MAX | 904.59 | 861.89 | 0.07 | 1.05 | 0.608 | 0.792 |
| MECOM | 812.03 | 546.49 | 0.57 | 1.49 | 0.041 | 0.228 |
| MEF2C | 3273.08 | 2056.23 | 0.67 | 1.59 | 0.008 | 0.123 |
| MKNK1 | 508.66 | 502.58 | 0.02 | 1.01 | 0.914 | 0.961 |
| MKNK2 | 2376.93 | 1777.66 | 0.42 | 1.34 | 0.019 | 0.169 |
| MOS | | | | | | |
| MRAS | 649.98 | 505.29 | 0.36 | 1.29 | 0.118 | 0.357 |
| MYC | 51.25 | 52.61 | -0.04 | 0.98 | 0.927 | 0.968 |
| NF1 | 983.49 | 763.84 | 0.36 | 1.29 | 0.139 | 0.381 |
| NFATC2 | 45.97 | 38.77 | 0.24 | 1.18 | 0.572 | 0.766 |
| NFATC4 | 594.88 | 679.88 | -0.19 | 0.88 | 0.376 | 0.615 |
| NFKB1 | 1088.91 | 1067.69 | 0.03 | 1.02 | 0.831 | 0.920 |
| NFKB2 | 294.06 | 301.69 | -0.03 | 0.98 | 0.832 | 0.920 |
| NGF | 29.22 | 44.69 | -0.60 | 0.66 | 0.250 | 0.504 |
| NLK | 319.53 | 394.98 | -0.30 | 0.81 | 0.184 | 0.433 |
| NR4A1 | 624.03 | 500.62 | 0.32 | 1.25 | 0.459 | 0.682 |
| NRAS | 959.53 | 857.89 | 0.16 | 1.12 | 0.387 | 0.624 |
| NTF3 | 1319.88 | 1249.19 | 0.08 | 1.06 | 0.681 | 0.835 |
| NTF4 | | | | | | |
| NTRK1 | | | | | | |
| NTRK2 | 1200.68 | 942.24 | 0.35 | 1.27 | 0.129 | 0.371 |
| PAK1 | 91.03 | 76.13 | 0.26 | 1.20 | 0.384 | 0.621 |

| Gene symbol | Mean Col3a1$^{+/G182R}$ (n=3) | Mean Col3a1$^{+/+}$ (n=4) | log2FoldChange | FoldChange | p-value | *adjusted p-value |
|---|---|---|---|---|---|---|
| PAK2 | 2163.01 | 1915.31 | 0.17 | 1.13 | 0.226 | 0.479 |
| PDGFA | 1693.88 | 2333.57 | -0.46 | 0.73 | 0.011 | 0.139 |
| PDGFB | 422.33 | 488.31 | -0.21 | 0.87 | 0.436 | 0.663 |
| PDGFRA | 1032.45 | 832.42 | 0.31 | 1.24 | 0.149 | 0.393 |
| PDGFRB | 5341.91 | 7036.08 | -0.40 | 0.76 | 0.016 | 0.160 |
| PLA2G10 | | | | | | |
| PLA2G12A | 478.52 | 507.66 | -0.08 | 0.94 | 0.486 | 0.703 |
| PLA2G12B | | | | | | |
| PLA2G1B | | | | | | |
| PLA2G2A | | | | | | |
| PLA2G2C | | | | | | |
| PLA2G2D | 6.96 | 8.65 | -0.31 | 0.81 | 0.665 | NA |
| PLA2G2E | 4.15 | 6.45 | -0.66 | 0.63 | 0.433 | NA |
| PLA2G2F | | | | | | |
| PLA2G3 | 7.54 | 16.69 | -1.13 | 0.46 | 0.081 | 0.304 |
| PLA2G4A | 1425.23 | 753.99 | 0.92 | 1.89 | 0.053 | 0.251 |
| PLA2G4B | 76.04 | 164.53 | -1.12 | 0.46 | 0.046 | 0.238 |
| PLA2G4E | 79.90 | 89.98 | -0.16 | 0.89 | 0.709 | 0.852 |
| PLA2G5 | 44.22 | 21.99 | 1.00 | 2.00 | 0.303 | 0.553 |
| PLA2G6 | 163.58 | 212.85 | -0.38 | 0.77 | 0.158 | 0.404 |
| PPM1A | 2924.94 | 2204.39 | 0.41 | 1.33 | 0.064 | 0.274 |
| PPM1B | 1814.08 | 1500.72 | 0.27 | 1.21 | 0.160 | 0.406 |
| PPP3CA | 906.09 | 628.29 | 0.53 | 1.44 | 0.006 | 0.112 |
| PPP3CB | 1309.42 | 1262.44 | 0.05 | 1.04 | 0.761 | 0.882 |
| PPP3CC | 285.65 | 237.62 | 0.26 | 1.20 | 0.086 | 0.310 |
| PPP3R1 | 1580.36 | 1081.13 | 0.55 | 1.46 | 0.049 | 0.245 |
| PPP3R2 | | | | | | |
| PPP5C | 741.91 | 870.32 | -0.23 | 0.85 | 0.058 | 0.262 |
| PRKACA | 1924.42 | 2017.08 | -0.07 | 0.95 | 0.707 | 0.851 |
| PRKACB | 3107.30 | 2316.73 | 0.42 | 1.34 | 0.126 | 0.367 |
| PRKACG | | | | | | |
| PRKCA | 620.36 | 479.87 | 0.37 | 1.29 | 0.172 | 0.420 |
| PRKCB | 10.48 | 16.04 | -0.63 | 0.65 | 0.221 | 0.473 |
| PRKCG | 149.48 | 263.86 | -0.82 | 0.57 | 0.014 | 0.151 |
| PRKX | 349.90 | 230.19 | 0.60 | 1.52 | 0.005 | 0.099 |
| PTPN5 | | | | | | |
| PTPN7 | 18.40 | 24.75 | -0.42 | 0.75 | 0.310 | 0.559 |
| PTPRR | 60.21 | 24.03 | 1.33 | 2.51 | 0.016 | 0.159 |
| RAC1 | 5543.21 | 4645.90 | 0.25 | 1.19 | 0.155 | 0.401 |
| RAC2 | 52.46 | 44.51 | 0.38 | 1.30 | 0.487 | 0.704 |
| RAC3 | 106.94 | 121.81 | -0.18 | 0.88 | 0.309 | 0.558 |
| RAF1 | 2585.25 | 3181.08 | -0.30 | 0.81 | 0.261 | 0.513 |
| RAP1A | 2475.75 | 1455.53 | 0.77 | 1.70 | 0.029 | 0.200 |
| RAP1B | 2387.84 | 1840.25 | 0.38 | 1.30 | 0.064 | 0.274 |
| RAPGEF2 | 933.16 | 737.49 | 0.34 | 1.26 | 0.158 | 0.404 |
| RASA1 | 662.43 | 641.42 | 0.05 | 1.03 | 0.797 | 0.902 |
| RASA2 | 680.62 | 488.31 | 0.48 | 1.39 | 0.026 | 0.192 |
| RASGRF1 | | | | | | |
| RASGRF2 | 227.60 | 160.13 | 0.51 | 1.42 | 0.149 | 0.393 |
| RASGRP1 | 9.41 | 4.15 | 1.11 | 2.15 | 0.228 | NA |
| RASGRP2 | 1168.22 | 1701.24 | -0.54 | 0.69 | 0.004 | 0.092 |
| RASGRP3 | 223.01 | 236.48 | -0.08 | 0.94 | 0.874 | 0.941 |
| RASGRP4 | 43.60 | 77.73 | -0.82 | 0.57 | 0.139 | 0.382 |
| RELA | 1249.30 | 1342.57 | -0.10 | 0.93 | 0.505 | 0.717 |
| RELB | 308.82 | 322.08 | -0.06 | 0.96 | 0.751 | 0.877 |
| RPS6KA1 | 56.33 | 43.71 | 0.38 | 1.30 | 0.421 | 0.651 |
| RPS6KA2 | 171.03 | 136.36 | 0.32 | 1.25 | 0.049 | 0.244 |
| RPS6KA3 | 1310.46 | 977.78 | 0.42 | 1.34 | 0.172 | 0.420 |
| RPS6KA4 | | | | | | |
| **RPS6KA5** | **349.16** | **170.58** | **1.03** | **2.04** | **1.170E-04** | **0.022** |
| RPS6KA6 | | | | | | |

| Gene symbol | Mean Col3a1$^{+/G182R}$ (n=3) | Mean Col3a1$^{+/+}$ (n=4) | log2FoldChange | FoldChange | p-value | *adjusted p-value |
|---|---|---|---|---|---|---|
| RRAS | 4597.08 | 5015.35 | -0.13 | 0.92 | 0.365 | 0.605 |
| RRAS2 | 901.30 | 778.94 | 0.21 | 1.16 | 0.392 | 0.628 |
| SOS1 | 534.48 | 408.06 | 0.39 | 1.31 | 0.048 | 0.242 |
| SOS2 | 469.58 | 391.58 | 0.26 | 1.20 | 0.268 | 0.521 |
| SRF | 994.80 | 674.22 | 0.56 | 1.47 | 0.087 | 0.311 |
| STK3 | 712.07 | 573.44 | 0.31 | 1.24 | 0.286 | 0.537 |
| STK4 | 731.37 | 610.12 | 0.26 | 1.20 | 0.306 | 0.555 |
| STMN1 | 300.42 | 421.67 | -0.49 | 0.71 | 0.114 | 0.350 |
| TAB1 | 325.96 | 500.98 | -0.62 | 0.65 | 0.005 | 0.105 |
| TAB2 | 1927.46 | 1533.83 | 0.33 | 1.26 | 0.096 | 0.326 |
| TAOK1 | 2371.50 | 1630.10 | 0.54 | 1.45 | 0.038 | 0.222 |
| TAOK2 | 1221.20 | 1552.45 | -0.34 | 0.79 | 0.131 | 0.372 |
| TAOK3 | 505.49 | 357.17 | 0.50 | 1.41 | 0.194 | 0.443 |
| TGFB1 | 361.51 | 348.43 | 0.06 | 1.04 | 0.830 | 0.920 |
| TGFB2 | 1660.69 | 1680.67 | -0.02 | 0.99 | 0.958 | 0.980 |
| TGFB3 | 3683.32 | 3348.98 | 0.14 | 1.10 | 0.444 | 0.670 |
| TGFBR1 | 2516.24 | 1794.57 | 0.49 | 1.40 | 0.140 | 0.382 |
| TGFBR2 | 3402.86 | 2293.28 | 0.57 | 1.48 | 0.043 | 0.232 |
| TNF | 4.65 | 4.85 | 0.01 | 1.00 | 0.994 | NA |
| TNFRSF1A | 1756.44 | 1831.48 | -0.06 | 0.96 | 0.581 | 0.773 |
| TP53 | | | | | | |
| TRAF2 | 493.05 | 508.31 | -0.04 | 0.97 | 0.809 | 0.908 |
| TRAF6 | 587.93 | 534.04 | 0.14 | 1.10 | 0.342 | 0.585 |

Significant differential expressed genes are mentioned in bold. Genes not found in transcriptomic data are highlighted in grey.
* Adjusted p-value with the Benjamini-Hochberg method.

**Supplemental Table 4 : Expression of genes of the PLC/IP3/PKC/ERK signaling pathway in the transcriptomic data of *Col3a1*$^{+/G182R}$**

| Gene symbol | IPA terms | *Col3a1* $^{+/G938D}$ | Mean *Col3a1* $^{+/G182R}$ (n=3) | Mean *Col3a1* $^{+/+}$ (n=4) | log2FoldChange | FoldChange | p-value | adjusted p-value |
|---|---|---|---|---|---|---|---|---|
| **Ldlr** | | Upregulated | **863.65** | **2078.79** | **-1.27** | **0.42** | **2.89E-05** | **0.011** |
| Serpine1 | | Upregulated | 5004.03 | 2534.79 | 0.98 | 1.97 | 6.04E-03 | 0.109 |
| Egr1 | | Upregulated | 1128.25 | 783.51 | 0.53 | 1.44 | 0.108 | 0.343 |
| F3 | | Upregulated | 1292.12 | 1123.77 | 0.20 | 1.15 | 0.370 | 0.609 |
| **Hspa1b** | | Upregulated | **1746.02** | **438.12** | **1.99** | **3.98** | **3.45E-06** | **0.005** |
| **Hspa1a** | | Upregulated | **2117.43** | **776.63** | **1.45** | **2.73** | **1.05E-04** | **0.021** |
| Fos | | Upregulated | 674.87 | 143.46 | 2.23 | 4.70 | 3.38E-03 | 0.089 |
| Bmp4 | | Upregulated | 457.50 | 363.78 | 0.33 | 1.26 | 0.337 | 0.582 |
| Jun | | Upregulated | 2036.19 | 1392.21 | 0.55 | 1.46 | 0.019 | 0.169 |
| Atf3 | | Upregulated | 713.27 | 300.27 | 1.25 | 2.38 | 1.32E-03 | 0.065 |
| Egr2 | | Upregulated | 17.70 | 14.09 | 0.23 | 1.17 | 0.669 | 0.828 |
| **Wwp2** | | Upregulated | **14224.97** | **26431.05** | **-0.89** | **0.54** | **1.27E-04** | **0.023** |
| Egr3 | | Upregulated | 38.39 | 36.16 | 0.13 | 1.09 | 0.779 | 0.891 |
| Id3 | | Upregulated | 4211.41 | 4664.02 | -0.15 | 0.90 | 0.443 | 0.669 |
| Bmf | | Upregulated | 660.42 | 1302.65 | -0.98 | 0.51 | 0.026 | 0.190 |
| Nuak1 | | Upregulated | 1600.10 | 1455.28 | 0.14 | 1.10 | 0.631 | 0.806 |
| Mfsd2a | | downregulated | 8.80 | 5.88 | 0.54 | 1.45 | 0.560 | NA |
| Apob | | downregulated | | | | | | |
| Tsc22d3 | | downregulated | 648.44 | 359.76 | 0.85 | 1.80 | 3.778E-03 | 0.092 |
| Zbtb16 | | downregulated | 1006.33 | 181.07 | 2.47 | 5.56 | 1.001E-03 | 0.059 |
| Ifi203 | | downregulated | 262.20 | 230.70 | 0.19 | 1.14 | 0.6966904 | 0.845 |
| Npvf | | unchanged | | | | | | |
| Polr2a | | unchanged | 749.15 | 707.18 | 0.08 | 1.06 | 0.679 | 0.834 |
| Msk1/2 | | unchanged | | | | | | |
| Cd3g | CD3 | unchanged | 4.75 | 9.46 | -0.95 | 0.52 | 0.304 | NA |
| Cd3d | | unchanged | 3.01 | 3.17 | -0.13 | 0.91 | 0.913 | NA |
| H3f3a | | unchanged | 11027.67 | 8503.92 | 0.37 | 1.30 | 0.103 | 0.336 |
| H3f3b | H3 | unchanged | 5704.62 | 6344.59 | -0.15 | 0.90 | 0.408 | 0.642 |
| H3f3c | | unchanged | 388.74 | 379.49 | 0.03 | 1.02 | 0.924 | 0.966 |
| Ccna2 | cyclin-a | unchanged | 88.11 | 143.34 | -0.70 | 0.62 | 0.106 | 0.340 |
| Ccna1 | | unchanged | | | | | | |
| | TCR | unchanged | | | | | | |
| | Ig | unchanged | | | | | | |
| Nfkb1 | | unchanged | 1088.91 | 1067.69 | 0.03 | 1.02 | 0.831 | 0.920 |
| Nfkb2 | | unchanged | 294.06 | 301.69 | -0.03 | 0.98 | 0.832 | 0.920 |
| Mapk8 (Jnk) | | Main effector | 688.91 | 488.57 | 0.49 | 1.41 | 0.052 | 0.249 |
| Mapk1 (Erk2) | | Main effector | 3174.48 | 2378.37 | 0.42 | 1.33 | 0.232 | 0.485 |
| Mapk3 (Erk1) | | Main effector | 2051.16 | 2105.50 | -0.04 | 0.97 | 0.757 | 0.880 |
| Mapk14 (p38 MAPK) | | Main effector | 1791.45 | 2057.68 | -0.20 | 0.87 | 0.052 | 0.249 |
| Akt1 | | Main effector | 3317.42 | 2918.34 | 0.18 | 1.14 | 0.313 | 0.561 |
| Akt2 | | Main effector | 3747.25 | 3344.54 | 0.16 | 1.12 | 0.220 | 0.472 |
| Akt3 | | Main effector | 2631.11 | 1878.04 | 0.49 | 1.40 | 0.091 | 0.317 |

Significant differential expressed genes are mentioned in bold. Genes not found in transcriptomic data are highlighted in grey.
Adjusted p-value with the Benjamini-Hochberg method.

**Supplemental Table 5 : Quantification of the 3 TGF-β isoforms in *Col3a1*$^{+/G182R}$ mice.**

| | Col3a1$^{+/+}$ (n=6) | Col3a1$^{+/G182R}$ (n=6) | p value |
|---|---|---|---|
| TGF-β1 | 34.4 ± 7.8 | 45.3 ± 3.8 | 0.240 |
| TGF-β2 | 29.7 ± 4.3 | 34.6 ± 3.9 | 0.420 |
| TGF-β3 | 10.4 ± 1.1 | 8.5 ± 1.3 | 0.288 |

Amount of the 3 TGF-β isoforms per µg of protein extracted from TA. Data are expressed as the mean ± SEM.
Student t-test.

**Supplemental Table 6 : Comparison of therapeutic strategies (monotherapies) with measurement of SBP and HR in *Col3a1*$^{+/G182R}$ male mice.**

| | Dose (mg/kg/j) / Age (weeks) | Propranolol Water | Propranolol Treatment 115 | Propranolol p value | Celiprolol Water | Celiprolol Treatment 250 | Celiprolol p value | Losartan Water | Losartan Treatment 135 | Losartan p value |
|---|---|---|---|---|---|---|---|---|---|---|
| SBP (mmHg) | 8 | 111.3 ± 2.5<br>n = 9 | 124.7 ± 2.4<br>n = 16 | **0.0016** | 118.3 ± 6.2<br>n = 7 | 112.2 ± 5.2<br>n = 6 | 0.4760 | 112.8 ± 3.6<br>n = 5 | 94.1 ± 3.1<br>n = 18 | **0.0067** |
| | 12 | 107.4 ± 3.0<br>n = 8 | 110.1 ± 2.5<br>n = 15 | 0.5147 | 116.4 ± 3.7<br>n = 9 | 109.9 ± 3.4<br>n = 4 | 0.3032 | 116.5 ± 5.6<br>n = 4 | 86.9 ± 3.6<br>n = 18 | **0.0015** |
| | 16 | 111.5 ± 3.4<br>n = 8 | 113.8 ± 4.2<br>n = 13 | 0.7102 | 119.4 ± 4.5<br>n = 11 | 121.7 ± 6.4<br>n = 5 | 0.7746 | 112.0 ± 2.9<br>n = 4 | 90.5 ± 4.4<br>n = 17 | **0.0326** |
| | 20 | 102.1 ± 4.1<br>n = 9 | 118.2 ± 3.6<br>n = 12 | **0.0082** | 115.0 ± 4.6<br>n = 11 | 113.5 ± 2.2<br>n = 9 | 0.7869 | 110.5 ± 9.6<br>n = 4 | 92.4 ± 6.1<br>n = 14 | 0.1695 |
| | 24 | 100.9 ± 0.8<br>n = 8 | 111.6 ± 5.3<br>n = 8 | 0.0662 | 114.0 ± 4.8<br>n = 11 | 125.5 ± 6.8<br>n = 8 | 0.1736 | 124.5 ± 8.5<br>n = 2 | 88.3 ± 3.4<br>n = 16 | **0.0023** |
| HR (bpm) | 8 | 641.2 ± 27.2<br>n = 9 | 446.7 ± 10.7<br>n = 16 | **<0.0001** | 605.3 ± 26.4<br>n = 7 | 550.5 ± 17.2<br>n = 6 | 0.1225 | 599.6 ± 18.4<br>n = 5 | 583.6 ± 18.5<br>n = 18 | 0.6684 |
| | 12 | 630.1 ± 21.9<br>n = 8 | 489.0 ± 10.6<br>n = 15 | **<0.0001** | 615.6 ± 28.3<br>n = 9 | 623.2 ± 21.7<br>n = 4 | 0.8700 | 581.0 ± 30.8<br>n = 4 | 641.8 ± 14.8<br>n = 18 | 0.0949 |
| | 16 | 650.6 ± 22.0<br>n = 8 | 460.8 ± 19.0<br>n = 13 | **<0.0001** | 635.3 ± 17.9<br>n = 11 | 651.3 ± 12.8<br>n = 5 | 0.5807 | 625.0 ± 22.0<br>n = 4 | 656.1 ± 15.2<br>n = 17 | 0.3641 |
| | 20 | 633.8 ± 32.1<br>n = 9 | 514.1 ± 25.1<br>n = 12 | **0.0077** | 633.5 ± 18.4<br>n = 11 | 635.7 ± 21.3<br>n = 9 | 0.9390 | 673.3 ± 19.4<br>n = 4 | 653.4 ± 17.9<br>n = 14 | 0.5825 |
| | 24 | 613.6 ± 39.8<br>n = 8 | 520.0 ± 23.6<br>n = 8 | 0.0626 | 654.4 ± 23.8<br>n = 11 | 661.9 ± 11.4<br>n = 8 | 0.8029 | 567.5 ± 90.5<br>n = 2 | 669.0 ± 10.8<br>n = 16 | **0.0204** |
| Mortality (%) | 24 | 46.7 | 40.0 | 0.6862 | 36.8 | 50.0 | 0.5411 | 62.5 | 17.6 | **0.0207** |

| | Dose (mg/kg/j) / Age (weeks) | Losartan at weaning Water | Losartan at weaning Treatment 135 | Losartan at weaning p value | Amlodipine 1%EtOH | Amlodipine Treatment 7.5 | Amlodipine p value | Hydralazine Water | Hydralazine Treatment 45 | Hydralazine p value |
|---|---|---|---|---|---|---|---|---|---|---|
| SBP (mmHg) | 8 | 112.8 ± 3.6<br>n = 5 | 107.8 ± 5.3<br>n = 13 | 0.5861 | 123.0 ± 3.5<br>n = 11 | 112.3 ± 2.8<br>n = 6 | 0.0603 | 122.0 ± 4.5<br>n = 18 | 120.4 ± 3.7<br>n = 15 | 0.7905 |
| | 12 | 116.5 ± 5.6<br>n = 4 | 96.5 ± 6.4<br>n = 10 | **0.0918** | 116.1 ± 3.6<br>n = 9 | 112.2 ± 3.2<br>n = 5 | 0.4818 | 118.5 ± 2.7<br>n = 18 | 112.6 ± 2.4<br>n = 13 | 0.1266 |
| | 16 | 112.0 ± 2.9<br>n = 4 | 105.5 ± 3.6<br>n = 11 | 0.3222 | 121.0 ± 6.8<br>n = 9 | 100.0 ± 7.2<br>n = 4 | 0.0914 | 122.7 ± 2.9<br>n = 19 | 114.9 ± 1.8<br>n = 13 | 0.0501 |
| | 20 | 110.5 ± 9.6<br>n = 4 | 98.0 ± 5.7<br>n = 3 | 0.3552 | 133.0 ± 4.0<br>n = 8 | 100.5 ± 3.3<br>n = 4 | **0.0004** | 117.6 ± 3.0<br>n = 19 | 114.3 ± 2.8<br>n = 12 | 0.4585 |
| | 24 | 124.5 ± 8.5<br>n = 2 | 80.2 ± 3.6<br>n = 11 | **0.0005** | 135.8 ± 5.2<br>n = 4 | 112.5 ± 8.3<br>n = 4 | 0.0548 | 117.3 ± 3.4<br>n = 17 | 113.3 ± 3.0<br>n = 12 | 0.4016 |
| HR (bpm) | 8 | 599.6 ± 18.4<br>n = 5 | 596.8 ± 30.6<br>n = 13 | 0.9580 | 632.8 ± 24.5<br>n = 11 | 702.7 ± 14.1<br>n = 6 | 0.0656 | 614.4 ± 17.7<br>n = 18 | 648.8 ± 23.1<br>n = 15 | 0.2393 |
| | 12 | 581.0 ± 30.8<br>n = 4 | 619.6 ± 18.9<br>n = 10 | 0.3001 | 662.3 ± 31.4<br>n = 9 | 673.2 ± 39.0<br>n = 5 | 0.8356 | 640.0 ± 15.9<br>n = 18 | 703.6 ± 12.8<br>n = 13 | **0.0065** |
| | 16 | 625.0 ± 22.0<br>n = 4 | 615.0 ± 25.0<br>n = 11 | 0.8244 | 673.6 ± 14.1<br>n = 9 | 756.0 ± 12.3<br>n = 4 | **0.0044** | 634.0 ± 14.1<br>n = 19 | 719.4 ± 9.2<br>n = 13 | **0.0001** |
| | 20 | 673.3 ± 19.4<br>n = 4 | 635.3 ± 50.7<br>n = 3 | 0.4677 | 717.5 ± 14.1<br>n = 8 | 786.8 ± 5.9<br>n = 4 | **0.0075** | 644.1 ± 12.0<br>n = 19 | 713.9 ± 8.8<br>n = 12 | **0.0002** |
| | 24 | 567.5 ± 90.5<br>n = 2 | 580.5 ± 31.6<br>n = 11 | 0.8771 | 732.0 ± 8.9<br>n = 4 | 748.0 ± 13.3<br>n = 4 | 0.3552 | 664.5 ± 15.9<br>n = 17 | 723.5 ± 8.1<br>n = 12 | **0.0070** |
| Mortality (%) | 24 | 62.5 | 7.7 | **0.0031** | 60.9 | 75.0 | 0.1261 | 50.0 | 29.9 | 0.1358 |

Data are expressed as the mean ± SEM. The number of mice at each time are indicated above the mean ± SEM.
Student-t test or Log-Rank test (Mantel-Cox) for the survival.

**Supplemental Table 7 : Comparison of therapeutic strategies (dual therapies in male mice and Amlodipine therapies in female mice) with measurement of SBP and HR in *Col3a1*$^{+/G182R}$ male mice.**

| | | Amlodipine-Propranolol males | | | Hydralazine-Celiprolol | | | Amlodipine females | | | Amlodipine-Propranolol females | | |
|---|---|---|---|---|---|---|---|---|---|---|---|---|---|
| | Dose (mg/kg/j) | Amlodipine 7.5 | Treatment A:7.5 P:115 | p value | Water | Treatment H:45 C:250 | p value | 1%EtOH | Treatment 7.5 | p value | Amlodipine 7.5 | Treatment A:7.5 P:115 | p value |
| | Age (weeks) | | | | | | | | | | | | |
| SBP (mmHg) | 8 | 112.3 ± 2.8 n = 6 | 99.8 ± 2.4 n = 5 | **0.0086** | 122.0 ± 4.5 n = 18 | 107.7 ± 5.6 n = 7 | 0.0900 | 118.3 ± 2.9 n = 18 | 103.7 ± 4.8 n = 3 | 0.0650 | 103.7 ± 4.8 n = 3 | 99.5 ± 2.4 n = 9 | 0.4186 |
| | 12 | 112.2 ± 3.2 n = 5 | 111.0 ± 1.5 n = 3 | 0.7937 | 118.5 ± 2.7 n = 18 | 105.0 ± 6.2 n = 9 | 0.0275 | 119.4 ± 2.7 n = 18 | 96.3 ± 9.5 n = 3 | **0.0070** | 96.3 ± 9.5 n = 3 | 102.8 ± 4.2 n = 9 | 0.4852 |
| | 16 | 100.0 ± 7.2 n = 4 | 89.5 ± 3.5 n = 2 | 0.3969 | 122.7 ± 2.9 n = 19 | 124.1 ± 3.7 n = 8 | 0.7955 | 122.8 ± 2.5 n = 17 | 100.5 ± 5.5 n = 2 | **0.0092** | 100.5 ± 5.5 n = 2 | 107.3 ± 5.4 n = 9 | 0.5886 |
| | 20 | 100.5 ± 3.3 n = 4 | 103.3 ± 5.5 n = 3 | 0.6572 | 117.6 ± 3.0 n = 19 | 106.0 ± 4.7 n = 8 | 0.0458 | 117.5 ± 3.1 n = 18 | 118.0 ± 8.0 n = 2 | 0.9610 | 118.0 ± 8.0 n = 2 | 89.4 ± 3.7 n = 8 | **0.0091** |
| | 24 | 112.5 ± 8.3 n = 4 | 89.0 ± 5.0 n = 2 | 0.1394 | 117.3 ± 3.4 n = 17 | 106.9 ± 4.4 n = 7 | 0.0944 | 118.3 ± 3.1 n = 13 | 117.0 ± 11.0 n = 2 | 0.8849 | 117.0 ± 11.0 n = 2 | 91.8 ± 4.7 n = 8 | **0.0467** |
| HR (bpm) | 8 | 702.7 ± 14.1 n = 6 | 518.4 ± 12.5 n = 5 | **<0.0001** | 614.4 ± 17.7 n = 18 | 608.2 ± 19.8 n = 7 | 0.8427 | 613.9 ± 14.4 n = 18 | 637.0 ± 27.2 n = 3 | 0.5412 | 637.0 ± 27.2 n = 3 | 516.2 ± 8.6 n = 9 | **0.0002** |
| | 12 | 673.2 ± 39.0 n = 5 | 616.7 ± 44.6 n = 3 | 0.3923 | 640.0 ± 15.9 n = 18 | 652.0 ± 6.0 n = 9 | 0.6062 | 648.6 ± 18.6 n = 18 | 690.7 ± 31.4 n = 3 | 0.3915 | 690.7 ± 31.4 n = 3 | 522.8 ± 25.7 n = 9 | **0.0063** |
| | 16 | 756.0 ± 12.3 n = 4 | 643.5 ± 35.5 n = 2 | **0.0168** | 634.0 ± 14.1 n = 19 | 653.3 ± 9.1 n = 8 | 0.4040 | 597.4 ± 30.8 n = 17 | 731.5 ± 10.5 n = 2 | 0.1639 | 731.5 ± 10.5 n = 2 | 513.5 ± 27.2 n = 9 | **0.0056** |
| | 20 | 786.8 ± 5.9 n = 4 | 621.0 ± 2.5 n = 3 | **<0.0001** | 644.1 ± 12.0 n = 19 | 673.2 ± 6.5 n = 8 | 0.1413 | 639.3 ± 22.8 n = 18 | 670.5 ± 90.5 n = 2 | 0.6763 | 670.5 ± 90.5 n = 2 | 518.1 ± 31.2 n = 8 | **0.0745** |
| | 24 | 748.0 ± 13.3 n = 4 | 625.0 ± 6.0 n = 2 | **0.0037** | 664.5 ± 15.9 n = 17 | 660.8 ± 10.3 n = 7 | 0.8870 | 646.6 ± 28.7 n = 13 | 742.0 ± 21.0 n = 2 | 0.2306 | 742.0 ± 21.0 n = 2 | 536.0 ± 22.9 n = 8 | **0.0028** |
| Mortality (%) | 24 | 75.0 | 70.0 | 0.3402 | 50.0 | 46.7 | 0.8477 | 25.0 | 50.0 | 0.1363 | 50.0 | 30.0 | 0.1365 |

Data are expressed as the mean ± SEM. The number of mice at each time are indicated above the mean ± SEM.
Student-t test or Log-Rank test (Mantel-Cox) for the survival.
